# Multi-modal tissue-aware graph neural network for *in silico* genetic discovery

**DOI:** 10.64898/2026.02.17.706433

**Authors:** Anusha Aggarwal, Ksenia Sokolova, Olga G. Troyanskaya

**Affiliations:** Quantitative and Computational Biology Program, Princeton University, NJ, USA; Lewis-Sigler Institute for Integrative Genomics, Princeton University, Princeton, NJ, USA; Princeton Precision Health, Princeton, NJ, USA; Center for Computational Biology, Flatiron Institute, New York, NY, USA; Department of Computer Science, Princeton University, Princeton, NJ, USA

## Abstract

Understanding how perturbations influence gene function in a tissue-specific manner is key to uncovering novel drug targets. However, current computational approaches emphasize global network or sequence-derived features over context-driven dependencies. We introduce Mahi, a scalable and interpretable graph neural network framework that learns gene representations by integrating chromatin accessibility, transcription factor binding, histone modifications, and protein structure features in tissue-specific contexts. Through pretraining on tissue-specific network topologies followed by multi-modal feature integration, Mahi learns context-aware gene embeddings across 290 tissues and cell-types. Mahi outperforms sequence-based models in predicting gene essentiality across 1,183 cancer cell lines, demonstrating the advantage of integrating molecular context and functional connectivity. The learned embedding space reveals tissue-specific functional organization, with genes forming distinct clusters reflecting their context-dependent roles. *In silico* gene knockout perturbations demonstrate Mahi’s ability to model intricate perturbation responses, identifying disease-relevant pathways and therapeutic targets. Together, these results demonstrate Mahi as a foundation for modeling tissue-specific gene function and perturbation responses, enabling applications in precision medicine, therapeutic target discovery, and prediction of context-dependent genetic vulnerabilities. All embeddings and the framework are publicly available to facilitate use by the scientific community.

## Introduction

A fundamental challenge in biology and precision medicine is understanding the effects of genetic perturbations. This is especially challenging considering seemingly simple genetic changes, such as gene knockouts, can have complex systems-level consequences and act differently in specific cellular and physiological contexts. These effects arise from complex networks of regulatory and biophysical processes that govern how genes and proteins are regulated and interact within specific cells or tissues. Accurately modeling genetic perturbations, therefore, requires integrating diverse modalities that capture transcriptional, post-transcriptional, structural, and interaction systems within their physiological context.

There has been substantial progress in learning gene and protein functions directly from sequence. Protein models such as ESM3^1^ and AlphaFold^2^ have pushed our ability to infer protein structure and function, while genomic sequence models like DeepSEA^3^, Sei^4^, and Enformer^5^ have uncovered extensive regulatory information encoded in the noncoding genome. Embeddings from these and other models^6–8^ have shown great promise for a diverse set of biomedical applications, but they capture each modality as a static entity without contextual integration. This is insufficient for accurate prediction of effects of genetic perturbations in context, nor for providing interpretability for these effects.

We introduce Mahi, a deep learning framework for multi-modal tissue- and cell-type-specific prediction of genetic perturbation effects. Mahi builds on multi-modal embeddings that leverage epigenetic and protein sequence models together with tissue-specific interaction maps, which are integrated through graph neural networks to model functional interactions across 290 tissues and cell types. We benchmark Mahi on a complex task of predicting the impact of single-gene knockouts, using binary tissue- and cell-type-specific essentiality across 1,183 human cancer cell lines, and show superior performance compared to genomic and protein sequence models. Beyond prediction, Mahi provides interpretable and biologically meaningful embeddings, where the learned embedding space reflects context-dependent functional organization with significantly higher dispersion across tissue-specific genes. Finally, Mahi supports *in silico* gene perturbation experiments, which produce interpretable tissue- and cell-type-resolved exploration of context-dependent network rewiring and disease-relevant pathways.

## Results

Mahi is a multi-modal, tissue- and cell-type-resolved graph neural network designed to model how genetic perturbations shape gene and protein function within their biological contexts. To capture shared and context-dependent relationships, we leverage network information for 19,073 protein-coding genes across 290 cell-type- and tissue-specific functional networks^9^, derived from large-scale gene expression data and protein-protein interactions (PPIs). We first pretrain an attention-based graph neural network (GNN) on a multigraph that integrates 35 core tissue-specific networks that are representative of the diversity of the full 290 tissue and cell-type contexts (Table S1).

We employ a self-supervised masked edge reconstruction task (ROCAUC=0.83, Figure S1) to learn generalizable network embeddings for each gene, capturing cross-tissue topology and shared interactions (Figure 1). These pretrained embeddings serve as shared initialization, providing a baseline of cross-tissue structure that is later refined within each of the 290 tissue and cell-type functional networks.

**Figure 1.**
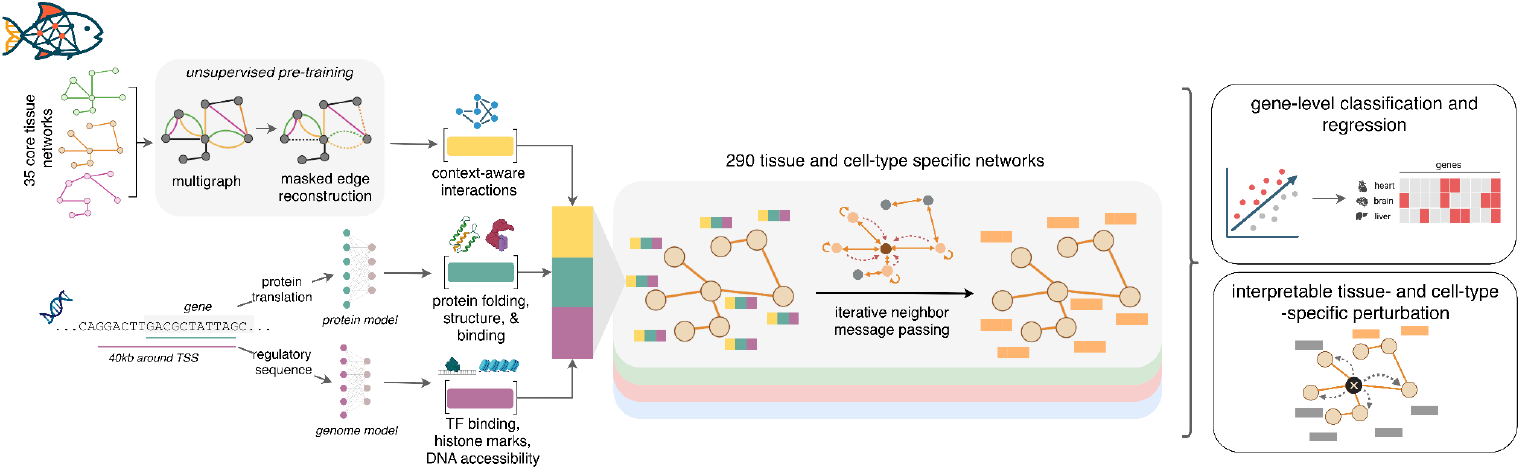
The Mahi framework integrates both molecular and network-level information. We construct a multigraph from 35 tissue-specific functional networks to capture gene interactions across diverse functional topologies, and learn context-aware embeddings via masked edge reconstruction. To complement these interaction-based features, we incorporate molecular signals from two domains: genomic information, which reflects how regulatory elements influence gene activity, and protein-level information, which captures biophysical and structural properties. By refining these signals across 290 tissue and cell-type networks, Mahi unifies regulatory, structural, and interaction modalities into a single functional representation, enabling gene-level prediction tasks and interpretable tissue- and cell-type-specific perturbation modeling.

To add functional context, we integrate molecular features that capture distinct biological aspects of gene function. Genomic embeddings^3^ describe the regulatory potential of each gene. Specifically, how chromatin accessibility, transcription factor binding, and histone modifications shape when and where a gene is expressed. Protein embeddings^1^, in contrast, encode how amino acid composition and structure determine a protein’s biochemical activity, stability, and interactions. Together, these representations link regulatory control with molecular action, providing complementary perspectives that Mahi integrates within tissue- and cell-type-specific networks.

To make the embeddings tissue- and cell-type specific, these modalities are aggregated and refined through iterative neighbor message passing, in which each gene’s representation is updated based on information from its network neighbors. This approach enables Mahi embeddings to capture both molecular and network-level information for each gene (Figure 1). These tissue and cell-type-specific embeddings serve as a foundation for downstream analyses, including prediction of gene essentiality, characterization of tissue-specific functional organization, and modeling of perturbation responses.

### Mahi accurately predicts cell-line-specific gene essentiality

We evaluated whether Mahi embeddings can predict context-specific gene essentiality across 1,183 human cancer cell lines from the Broad Institute’s DepMap^10^ project. DepMap provides genome-wide CRISPRi screen data measuring the effect of gene knockout on cell viability (Figure 2A). In each cell line, only about 10%-12% of genes are classified as essential, resulting in a highly imbalanced essentiality profile and making this a challenging prediction task.

**Figure 2.**
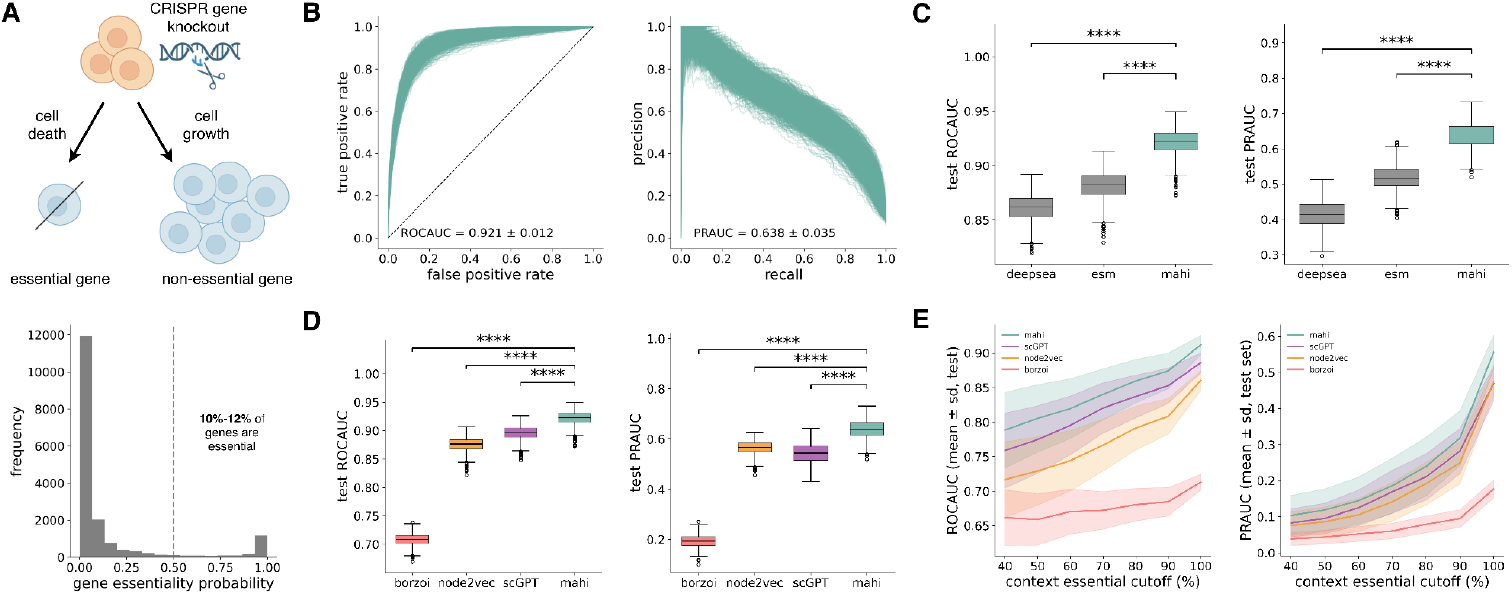
Mahi outperforms baselines in gene essentiality prediction. (A) Overview of gene essentiality data collection by the DepMap project, which measures cell growth following CRISPRi knockouts as a proxy for gene essentiality. The distribution of scores in the CaSki cell line indicates that approximately 10-12% of genes are essential. (B) Test set performance of Mahi on gene essentiality prediction across 1,183 cell lines. Mahi predicts essentiality with a high average ROCAUC of 0.921 and average PRAUC of 0.638. (C) Mahi embeddings achieve higher performance than the individual input features of its framework, emphasizing the need to consider all diverse modalities for biological prediction tasks. Statistical significance was assessed using two-sided paired t-tests (see Methods). (D) Mahi outperforms other sequence-, expression-, and graph-based embeddings in predicting gene essentiality. Statistical significance was assessed using two-sided paired t-tests (see Methods). (E) Comparison of test set performance on context-essential genes defined at varying frequency thresholds. Thresholds represent the maximum fraction of cell lines in which a gene is labeled as essential (e.g., <40%). As the threshold relaxes, predictive performance increases, but Mahi consistently outperforms scGPT, node2vec, and Borzoi embeddings across all thresholds.

For each cell line, we trained independent gradient-boosted tree models using the corresponding tissue-specific Mahi embeddings (Table S2). Model performance was evaluated on a held-out test set, with additional cross-validation analyses (see Methods). Mahi achieves consistently strong predictive accuracy across diverse cancer cell lines (test ROCAUC = 0.921 ± 0.012, PRAUC = 0.638 ± 0.035, CV ROCAUC = 0.917 ± 0.011, PRAUC = 0.638 ± 0.031; Figure 2B, Figure S2), demonstrating robust generalization across training folds and held-out gene sets.

### Mahi outperforms sequence-, expression-, and graph-based embeddings

To assess the contribution of our model’s input feature modalities, we compared Mahi to protein-level embeddings from ESM-C and sequence-based regulatory embeddings from DeepSEA. Mahi significantly outperformed the sequence-based embeddings across cell lines, demonstrating the power of integrating molecular features with tissue-specific network context (Figure 2C). We further compared Mahi to a set of sequence-, expression-, and graph-based representations. For expression-related baselines, we included Borzoi^11^, a sequence-to-expression deep learning model, as a strong external baseline, given that gene expression has been reported as a top predictive feature of gene essentiality^12,13^. We also included scGPT^8^, whose gene embeddings reflect large-scale expression variation across tissues and cell states. Since Mahi incorporates graph structure, we also compared it against node2vec^7^, a state-of-the-art algorithm to produce graph-based embeddings. Mahi significantly outperformed all sequence-, expression-, and graph-based approaches (Figure 2D), underscoring the value of integrating molecular features with tissue-specific network context.

To further evaluate Mahi’s ability to capture context-dependent essentiality, we examined its performance on genes that display variable essentiality across cell lines (“context essential” genes). Unlike universally essential (“pan-essential”) genes, context essential genes exhibit cell-type- or tissue-specific dependencies that are often harder to predict. The distinction between these categories is not absolute, as it depends on the number and diversity of the profiled cell lines. Therefore, we assessed performance across a range of thresholds that define how frequently a gene is essential (see Methods). Across all thresholds, Mahi consistently outperformed sequence-, expression-, and graph-based representations, underscoring its capacity to model both pan- and context-specific determinants of essentiality (Figure 2D).

### Mahi embeddings capture tissue-specific gene functionality

To assess whether Mahi embeddings capture biologically meaningful tissue-specific signals, we visualized the gene representations for each of the 290 tissues and cell-types using principal component analysis (PCA). The resulting two-dimensional projections revealed distinct clustering patterns across tissues, with tissue-specific genes displaying broader spatial separation than housekeeping genes, consistent with the higher variability of genes with tissue-specific functions (Figure 3A).

**Figure 3.**
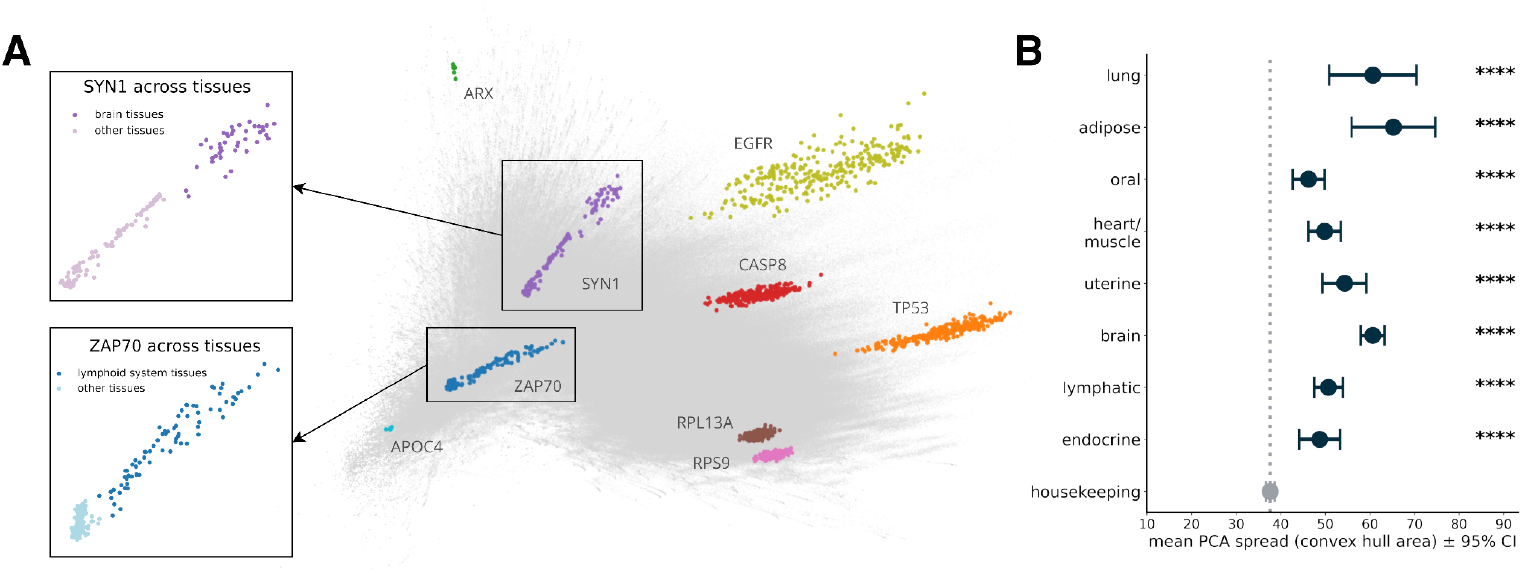
Mahi embeddings capture and distinguish tissue-specific gene function. (A) Principal component analysis (PCA) of protein-coding gene embeddings across 290 tissues reveals distinct tissue-specific clustering patterns. Each point represents the embedding of a gene in a specific tissue, with nine representative genes highlighted by color. The embedding distribution of SYN1 shows a clear outlier cluster corresponding to brain-associated tissues, consistent with its established role in neurotransmission and synaptic development. ZAP70, an immune response gene, exhibits an outlier cluster in lymphoid tissues, reflecting its tissue-specific expression profile. (B) Quantification of embedding dispersion across tissues for housekeeping genes and tissue-elevated genes defined by the Human Protein Atlas (HPA). The spread of points in PC space was quantified using convex hull area, with 95% bootstrap confidence intervals shown as error bars. Tissue-elevated genes exhibit significantly larger spread, reflecting tissue-specific distribution patterns, whereas housekeeping genes show reduced spread consistent with ubiquitous expression. Statistical significance was determined from the bootstrap distribution of mean differences (p-value < 1e-4 for all depicted tissues; see Methods).

We quantified the extent of tissue-specific variation by measuring how widely each gene’s embeddings are distributed in PCA space, using a convex hull-based measurement (see Methods). Convex hull areas were significantly larger for genes annotated as tissue-elevated by the Human Protein Atlas^14^ (HPA) than housekeeping genes (p-value < 1e-4 for all depicted tissues; Figure 3B, see Methods), confirming that Mahi representations systematically encode contextual diversity for tissue-specific genes.

Inspection of individual genes highlighted biologically coherent patterns: for example, SYN1, a neurotransmission-related gene involved in synaptic development^15^, localized to clusters corresponding to brain and neural tissues (e.g., brain, cerebellum, spinal cord); whereas ZAP70, an immune signaling gene^16^, appeared as an outlier with lymphoid and hematopoietic tissue clusters (e.g., lymphocyte, t-cells, blood). Conversely, ribosomal subunit genes such as RPLP0 and RPS9, which are housekeeping genes expressed in nearly all tissues^17,18^, displayed tightly compact clusters, reflecting their stable and ubiquitous functional roles.

### Mahi provides an interpretable framework for modeling context-specific perturbation responses

To evaluate whether Mahi embeddings can model the downstream effects of gene perturbations, we simulated *in silico* knockouts (KOs) by removing a single target gene and its direct edges from the relevant network and re-evaluating the pretrained model on the perturbed topology (Figure 4A). We quantified the effect of the perturbation by computing the Euclidean distance between the wild-type and perturbed embeddings for every gene in the network. These distances were normalized to the average distance obtained from 200 random single-gene KOs in the global network by calculating fold change for each gene (see Methods). The top 1,000 genes with the largest fold change for each target gene KO were then analyzed for Gene Ontology Biological Process enrichment using the PANTHER^19^ (Pan-GO) overrepresentation test.

**Figure 4.**
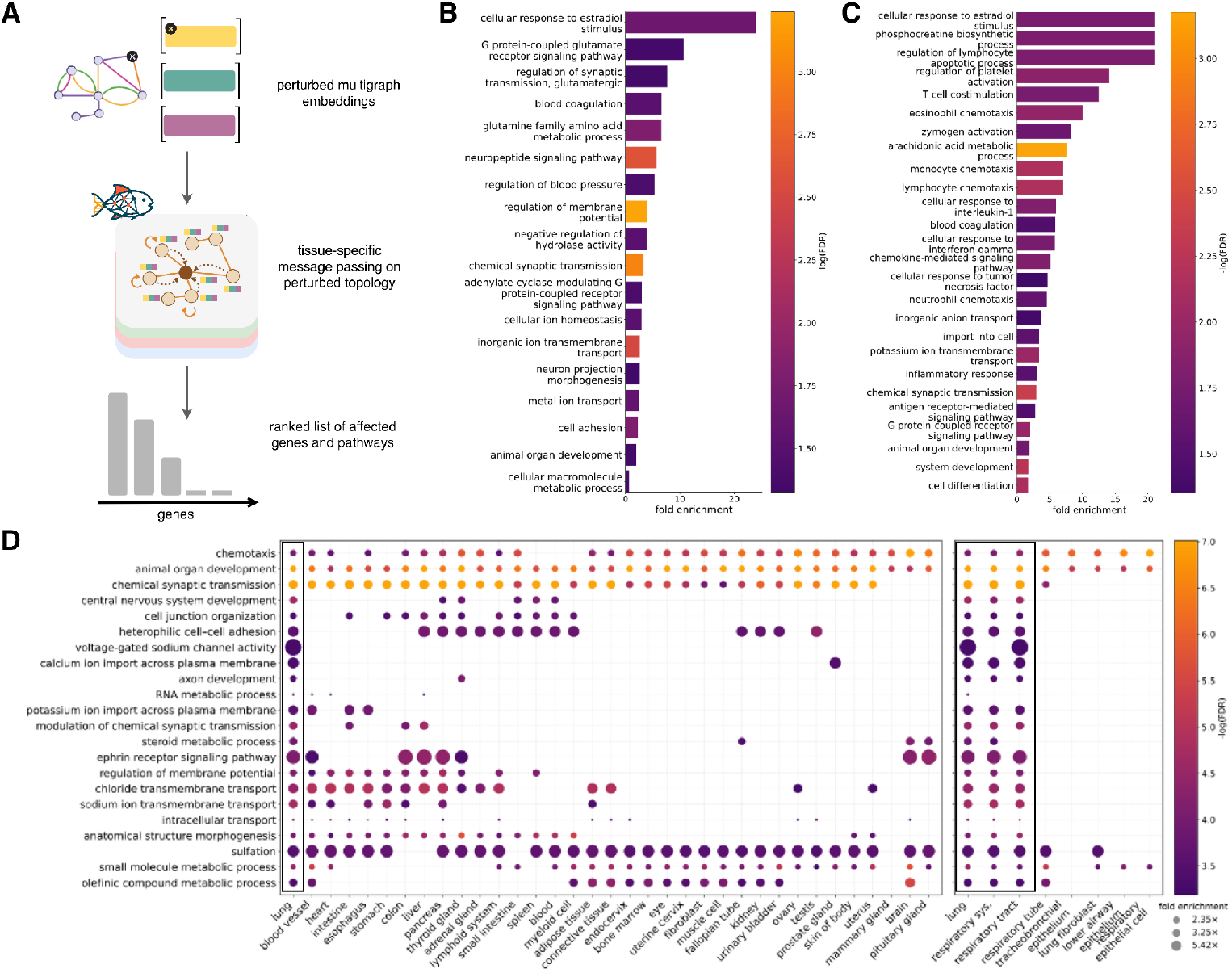
Mahi models context-specific gene perturbations. (A) Overview of the perturbation framework. To simulate a gene knockout, the target gene and its associated edges are removed from the multigraph, node embeddings are recomputed, and the same gene is also removed from tissue-specific message passing. (B) Pan-GO biological processes (BP) for ALPK3 knockout in heart tissue, simulating cardiomyopathy. Top enriched terms include circulatory system processes. (C) Pan-GO BP for DMD knockout in skeletal muscle tissue, simulating Duchenne muscular dystrophy. Top enriched terms include inflammatory and immune signals. (D) Pan-GO biological processes for CFTR knockout in lung tissue, compared to enrichment in 34 other core tissues (left) and additional tissues and cell-types from the full set of networks (right). Different pathways are enriched across the core tissues, whereas related tissues depict similar enrichment profiles.

We first examined *in silico* knockout of ALPK3, a kinase gene implicated in cardiomyopathy according to the OMIM^20^ database. The top perturbed genes following ALPK3 knockout in the heart tissue were enriched for processes involved in circulatory system function, such as blood coagulation, regulation of blood pressure, and homeostasis, consistent with core physiological dysregulation of cardiomyopathy (Figure 4B). Notably, enrichment of coagulation aligns with treatment protocols where anticoagulant therapies are used to reduce heart failure^21,22^. Beyond these canonical pathways, enrichment of ion transport, hormone signaling, and G-coupled protein receptor processes suggests secondary effects to cardiac regulation, which are also perturbed in heart failure and cardiomyopathy^23–25^.

Next, we analyzed DMD, a gene associated with Duchenne muscular dystrophy (DMD). The top perturbed genes were strongly enriched for immune and inflammatory processes, such as chemokine-mediated signaling, lymphocyte and monocyte chemotaxis, and responses to TNF, IL-1, and interferon, reflecting the chronic inflammation and immune cell infiltration that drive muscle necrosis in DMD^26^ (Figure 4C). Enrichment of the arachidonic acid metabolic process also reflects these inflammatory signatures and aligns with the use of anti-inflammatory therapies in its treatment^27^. Other phosphocreatine biosynthetic and ion transport processes were enriched, consistent with altered energy metabolism and disrupted membrane excitability related to muscular degeneration^28,29^.

### Mahi reveals how genetic perturbations drive distinct, tissue-specific pathways across context

Mahi can resolve how gene perturbations propagate through different tissues to produce distinct, pathway-specific mechanisms. To illustrate this, we perturbed CFTR, a gene encoding an ATP-gated chloride channel whose dysfunction causes cystic fibrosis, across multiple tissues. As expected, the lung network exhibited strong enrichment for ion transport and epithelial organization processes, including the chloride and sodium ion transmembrane transport, regulation of voltage-gated sodium channel activity, and cell junction organization (Figure 4D). These processes mirror the disrupted ion homeostasis and compromised epithelial barrier function that characterizes cystic fibrosis, highlighting CFTR’s role in maintaining airway hydration and mucosal clearance^30,31^.

When we applied the same perturbation across 34 other core tissues, the resulting pathway profiles varied substantially, highlighting tissue-specific regulatory programs rather than a uniform response. In contrast, lung-adjacent contexts such as the respiratory system and respiratory tract exhibited enrichment patterns highly concordant with the lungs, consistent with shared epithelial structure and regulatory architecture among related tissues. Strikingly, Mahi also uncovered biology far beyond canonical CFTR function in lung tissue. Several reproductive tissues, including the ovary, fallopian tubes, uterus, cervix, and testis, displayed coherent pathway responses to CFTR perturbation that aligned with known but subtle roles of CFTR in fertility (Figure S3). In the fallopian tube, for example, Mahi highlighted inflammatory and fluid-regulatory programs mirroring the epithelial inflammation and aberrant fluid secretion that drive hydrosalpinx-associated infertility^32^. Similarly, CFTR perturbations in testis-adjacent contexts showed activation of development and immune pathways consistent with congenital absence of the vas deferens in cystic fibrosis patients^33^. In cervical and endocervical tissues, Mahi recovered pathways central to hormone-dependent regulation of CFTR and its role in epithelial hydration^34^. These findings demonstrate that Mahi captures distinct, tissue-embedded regulatory programs across both widely-recognized affected tissues and those with more nuanced, tissue-specific phenotypic effects.

## Discussion

Mahi integrates molecular and structural gene and protein features with tissue-specific networks to enable *in silico* genetic analyses in cellular and tissue context. Mahi’s ability to simulate gene knockouts and recover relevant biological pathways highlights its potential for interpretable modeling of genetic perturbations. For example, we show that Mahi can predict tissue-dependent consequences of CFTR perturbation, revealing distinct mechanistic pathways that may guide tissue-specific therapeutic strategies. This can enable new opportunities in drug discovery and therapeutic hypothesis generation, where *in silico* perturbation of target genes within specific tissue networks could reveal downstream functional consequences and candidate pathways for intervention.

Overall, Mahi represents a step toward predictive and interpretable models of gene function in context. By embedding genes within their molecular and tissue-specific neighborhoods, it provides a scalable framework for understanding genetic variation, perturbation, and tissue environment. The learned embeddings recapitulate known tissue-specific organization and disease-relevant network responses and our findings demonstrate that incorporating tissue-specific network topology yields more biologically realistic representations of gene function than sequence alone. Since Mahi operates directly on multi-modal molecular features, it can also be extended to incorporate sequence variation or transcript isoform differences. Looking forward, Mahi could be expanded for perturbation modeling across a broader range of biological contexts, including development, aging, and other modalities. Mahi is readily adaptable to new data types and biological contexts, enabling richer and context-aware representations of gene function. The Mahi framework and precomputed embeddings are available on GitHub (https://github.com/FunctionLab/mahi).

## Methods

### Data collection & processing

The hg38.p14 gene annotation file was downloaded from NCBI^35^ and filtered for protein-coding genes with MANE (canonical) transcripts containing annotated TSSs, start/stop codons, and CDS regions. Functional networks were processed by retaining edges among protein-coding genes, ranking by weight, and keeping the top 3%, after which 35 tissue-specific networks (Table S1) were aggregated into a multigraph. CRISPR gene dependency scores for 1,183 cancer cell lines were taken from DepMap Public 25Q2, with dependency probabilities ≥ 0. 5 labeled as essential and < 0. 5 as non-essential per DepMap’s recommendations. Tissue-elevated and housekeeping gene sets were downloaded from Human Protein Atlas (HPA) version 24.1, and tissue-elevated sets were grouped into broader categories using HPA’s hierarchical tissue classification.

### Mahi Framework

#### Generating per-gene sequence embeddings

Genomic representations were generated by extracting 41,800 bps centered around the TSS of each gene as annotated in NCBI’s gene annotation file. Sequences were fed into DeepSEA in overlapping 2,000 bp windows, sliding every 200 bp, resulting in 2002 chromatin features per window. Forward and reverse strand embeddings were averaged and exponential decay weights were applied to emphasize widows closest to the TSS. The resulting genomic embedding is 1 x 2002 for every gene.

Protein representations were generated by extracting the start/stop codons and CDS regions for MANE (canonical) transcripts from NCBI’s gene annotation file. Codons were concatenated and translated to amino acid sequences using `Bio.seq`. Genes with invalid protein sequences (e.g., missing start codon “M” or lacking in-frame stop codon) were removed. The amino acid sequences were fed into ESM-C (600M), and residue embeddings were averaged to produce 1 x 1152 vector per gene.

#### Generating per-gene multigraph embeddings

A multi-graph *G* = (*V, E*) was constructed over genes *V*, with undirected, tissue-specific edges *E*. Each edge was one-hot encoded over 35 tissues. Node features were randomly initialized with 512-dimensional embeddings. Gene representations were trained using self-supervised masked edge reconstruction: 15% of edges in each mini-batch were masked, and the model used the remaining graph structure to predict the missing edge embeddings. The architecture comprised of *L* = 4 TransformerConv layers, followed by a small feed-forward neural network for edge prediction. Hyperparameters (learning rate, hidden dimensions, attention heads, dropout, and others) were optimized via Bayesian search in Weights & Biases^36^. A single TransformerConv layer can be modeled as:

#### Projections

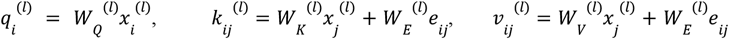

#### Attention

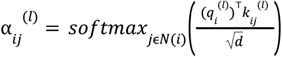

#### Aggregation

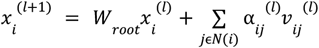

Here, *x*_*i*_ is the input node feature, *e*_*ij*_ is the edge feature between nodes *i* and *j*. *W*_*Q*_, *W*_*K*_, *W*_*V*_ generate the query, key, and value vectors *q, k, v* representations and *W*_*E*_ projects edge features. *N*(*i*) denotes the neighbors of node *i. W*_*root*_ is the root weight. The node embedding size *d* is fixed at 512 for all layers. α_*ij*_ is the attention coefficient controlling how much information node *i* receives from *j*. Each TransformerConv layer uses LayerNorm, ReLU, dropout and residual connections.

For masked edges (*i, j*), we concatenate node embeddings and pass them through a two-layer MLP with ReLU and dropout to predict tissue-wise logits. The MLP is defined as:

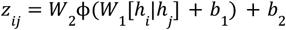

where *h*_*i*_ is the final node embedding after *L* TransformerConv layers 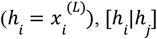 is the concatenated node embeddings, ϕ is ReLU, and *z*_*ij*_ is the predicted edge embedding. *W*_1_, *W*_2_ and *b*_1_, *b*_2_ are the MLP weights and biases.

We treat tissue reconstruction as a multi-label classification with a binary target per tissue. To train the model, binary cross-entropy loss was used for learning. The loss function for a single masked edge (*i, j*) is:

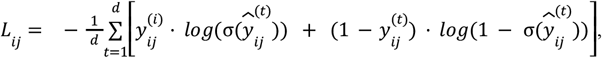

where *d* = 35 tissues, 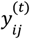 is the true binary label for tissue *t*, and 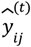 is the logit predicted by the model. σ is the sigmoid function. The final loss (over batch) is calculated as an average over a masked set of edges *M*,

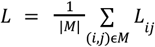

Nodes were split into 70% train, 10% validation, and 20% test. To avoid data leakage, only edges within each split were retained. Training used subgraph mini-batching, sampling up to 20 neighbors per node over 4 hops. The model was trained over 20 epochs with the Adam optimizer^37^ (learning rate = 0.001, batch size = 128). The encoder produces a 512-dimensional embedding per gene.

#### Integrating embeddings with graph-based propagation for tissue-specificity

Sequence and graph embeddings were concatenated and refined with APPNP^38^, a propagation-only method that diffuses features over tissue-specific networks. APPNP is defined as:

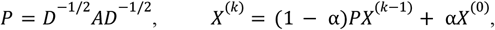

where *A* is the tissue adjacency matrix (with self-loops added), *D* its degree matrix, *P* the normalized adjacency, *X* the concatenated node features, *k* the propagation step, and α the teleport probability. We performed a grid search over *k* ϵ {1, 2, 3, 4, 5, 6, 7, 8, 9, 10, 15, 20, 25, 30} and α ϵ {0. 1,…, 0. 9}, selecting *k* = 8 and α = 0. 6 as the optimal balance of neighbor and original features.

We use APPNP to make the pretrained multi-modal embeddings context-specific by propagating them over tissue- and cell-type-specific networks, allowing for consistent control of message passing without oversmoothing the embeddings.

### Predicting downstream functional tasks & analysis of embedding space

#### Gene essentiality prediction across all cancer cell lines

For each of the 1,183 cell lines, we trained a separate XGBoost^39^ model to predict gene essentiality using corresponding tissue-specific Mahi embeddings. Each cancer cell line was mapped to 1 of 290 tissues based on its annotated cancer type and OncoTree lineage in DepMap (Table S2). Genes were randomly split into 80% training and 20% testing, with 5-fold stratified cross-validation within the training set to tune performance. Mean ROCAUC and PRAUC across folds were recorded, and final performance was evaluated on the held-out 20% using the model from the last fold. XGBoost models used a learning rate of 0.05, max tree depth of 4, 400 trees, and a subsample ratio of 0.8.

Borzoi embeddings were generated using the Replicate 0 (Human) checkpoint. Per-gene vectors taken from the encoder output layer (*add_15*) with global average pooling. Genomic sequences of 524,288 bps around the TSS were one-hot encoded and processed with the released Borzoi model, corresponding to the maximum input sequence length supported by Borzoi and the authors’ recommended setting for capturing long-range regulatory interactions. scGPT and node2vec embeddings were obtained from a prior benchmarking study^6^, and were filtered to match Mahi’s gene set. Differences in metrics between Mahi and baseline models (DeepSEA, ESM-C, scGPT, node2vec and Borzoi) were evaluated using two-sided paired t-tests across the 1,183 cell lines.

To assess performance across varying essentiality levels, genes were stratified by the fraction of cell lines in which they were essential. Since the notion of context essentiality depends on cell line coverage, we evaluated thresholds *t* ϵ {0. 4, 0. 5, 0. 6, 0. 7, 0. 8, 0. 9, 1. 0}. For each threshold, the evaluation set included genes essential in fewer than *t* × 100% of the profiled cell lines (with frequency ≥ 0 and < *t*).

#### Clustering analysis

To analyze tissue-specific structure in the embedding space, we performed principal component analysis (PCA) on standardized gene-tissue embeddings from all tissue-specific networks. We focused on the first two components, which capture a substantial fraction of the variance in the embedding space, for both visualization and quantitative analysis. Tissue separation was measured by computing the convex hull area using SciPy’s^40^ convex hull function (with `qhull_options =` ‘`QJ`’). Hull areas for tissue-enhanced and housekeeping genes were compared to assess tissue-specific dispersion.

Statistical significance was evaluated via bootstrap resampling: 1,000 bootstrap samples of mean hull area were generated for each group, and a one-sided p-value was obtained from the proportion of resampled mean differences ≤ 0. 95% confidence intervals were derived from the bootstrap percentiles.

#### Perturbation analysis

*In silico* knockouts (KOs) were simulated by removing a target gene and its direct edges from the tissue-specific graph. The pretrained GNN was then run on the perturbed topology to produce updated embeddings, reflecting the network’s response to gene loss. During tissue-specific propagation, the knocked-out gene and its edges were also removed from the corresponding tissue-specific network.

For each gene, the perturbation-induced shift in functional representation was quantified as the Euclidean distance *d* between the wild-type (*E*_*WT*_) and perturbed embeddings (*E*_*KO*_):

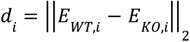

To account for baseline network sensitivity, we generated a background distribution by performing 200 random KOs in the global network and computing their distances. The normalized perturbation score for each gene was defined as the fold change between its distance and the mean random distance:

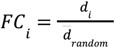

Genes were ranked by *FC*_*i*_, and the top 1,000 genes were used for enrichment analyses. GO Biological Process terms were identified using PANTHER (Pan-GO) overrepresentation test (based on Fisher’s exact test). Background distances were estimated using increasing numbers of random gene knockouts, with 200 knockouts used as a conservative choice after initial stabilization of gene rankings.

## Acknowledgements

We thank all members of the Troyanskaya lab for their constructive feedback and discussions during the preparation of this manuscript. This work was conducted utilizing the computing resources, supported by the Scientific Computing Core, at the Flatiron Institute. This work was supported by funding from the National Institutes of Health (NIH) under grants U01DK133090 and 2U24DK100845-12 and the Bill & Melinda Gates Foundation under grant INV-081342.

Figures 1 and parts of Figures 4 were generated using BioRender.com.

## Author Contributions

A.A., K.S., and O.G.T. conceived the study. A.A. and K.S. designed the study, developed the computational methods and performed the analyses. K.S. and O.G.T. supervised the project and contributed to its conceptual development. A.A., K.S., and O.G.T. wrote the manuscript. All authors reviewed and approved the final manuscript.

## Data availability

Downloadable links from HumanBase for the functional networks can be found in Table S3. Gene essentiality data was obtained from DepMap Public 25Q2 release. Tissue-elevated and housekeeping gene sets were downloaded from Human Protein Atlas (HPA) version 24.1.

## Code availability

The Mahi framework code is available at https://github.com/FunctionLab/mahi and the model and associated data files can be downloaded by following the instructions in the GitHub repository.

## Supplement

**Table S1**. Core 35 tissues used for training the multigraph on self-supervised masked edge reconstruction task.

**Table S2**. Mapping of Mahi embeddings to each of the 1,183 cell lines according to the cancer type and OncoTree lineage annotations provided by DepMap.

**Table S3**. Downloadable links from HumanBase for each of the tissue- and cell-type-specific functional networks.

**Figure S1.**
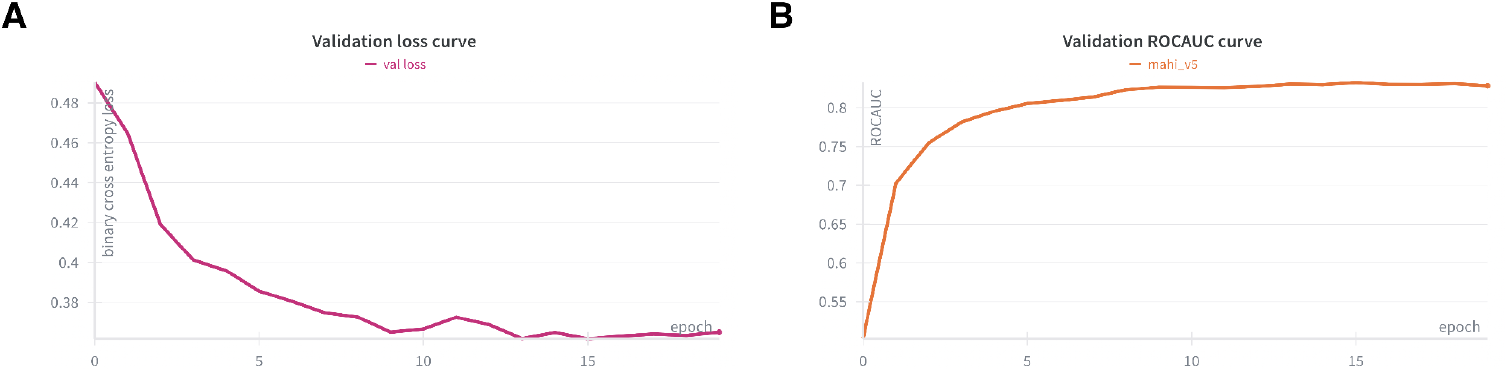
Validation curves for multigraph masked edge reconstruction. (A) Validation loss curve of training task using binary cross entropy. Validation loss converges at 0.37 after 20 epochs. (B) ROCAUC curve of validation set on the binary prediction across 35 tissues. ROCAUC converges at 0.83 after 20 epochs.

**Figure S2.**
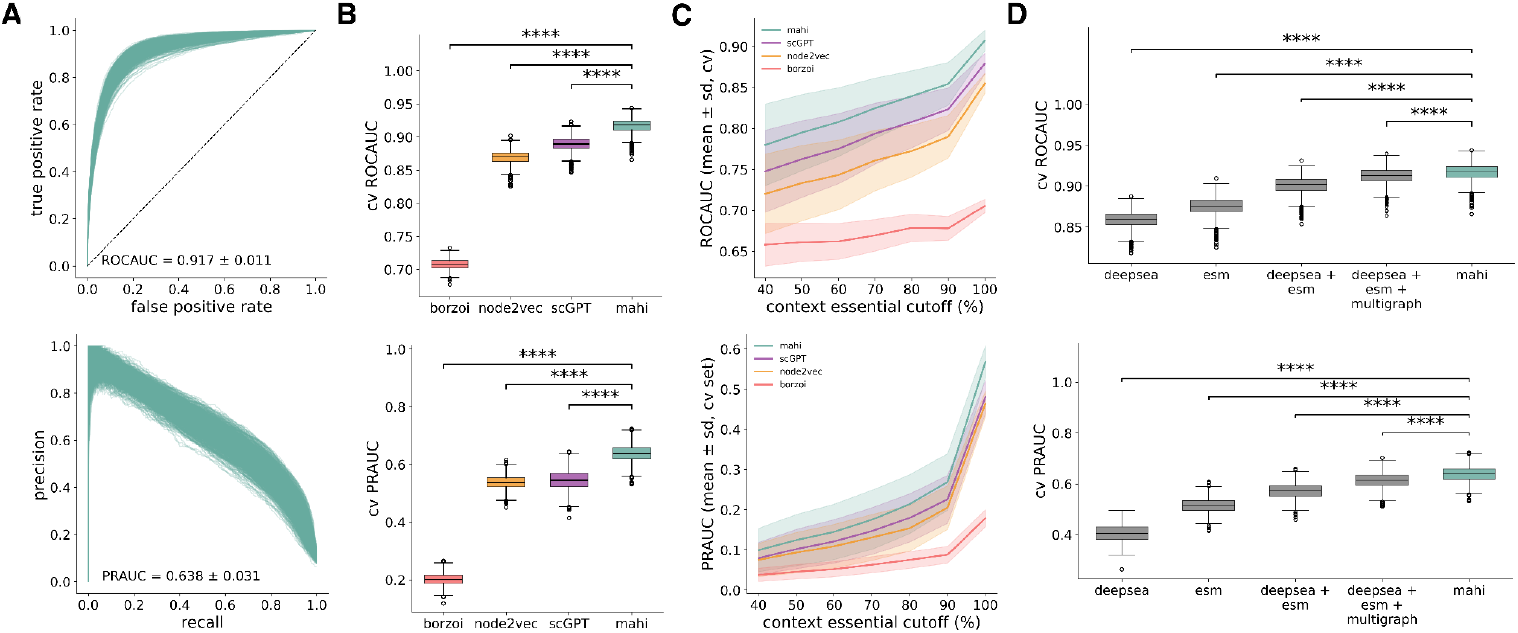
Cross validation performance of gene essentiality prediction task. (A) Cross validation (CV) performance of Mahi on gene essentiality prediction across 1,183 cell lines. Mahi predicts essentiality with a high average ROCAUC of 0.917 and average PRAUC of 0.638. (B) Mahi outperforms other sequence-, expression, and graph-based embeddings in predicting gene essentiality. Statistical significance was assessed using paired t-tests (see Methods). (C) Comparison of CV performance on context-essential genes defined at varying frequency thresholds. Thresholds represent the maximum fraction of cell lines in which a gene is labeled as essential (e.g., <40%). As the threshold relaxes, predictive performance increases, but Mahi consistently outperforms scGPT, node2vec, and Borzoi embeddings across all thresholds. (D) Mahi ablation study comparing the individual components of the framework and their concatenated (ablated) counterparts. Mahi consistently outperforms all ablated variants.

**Figure S3.**
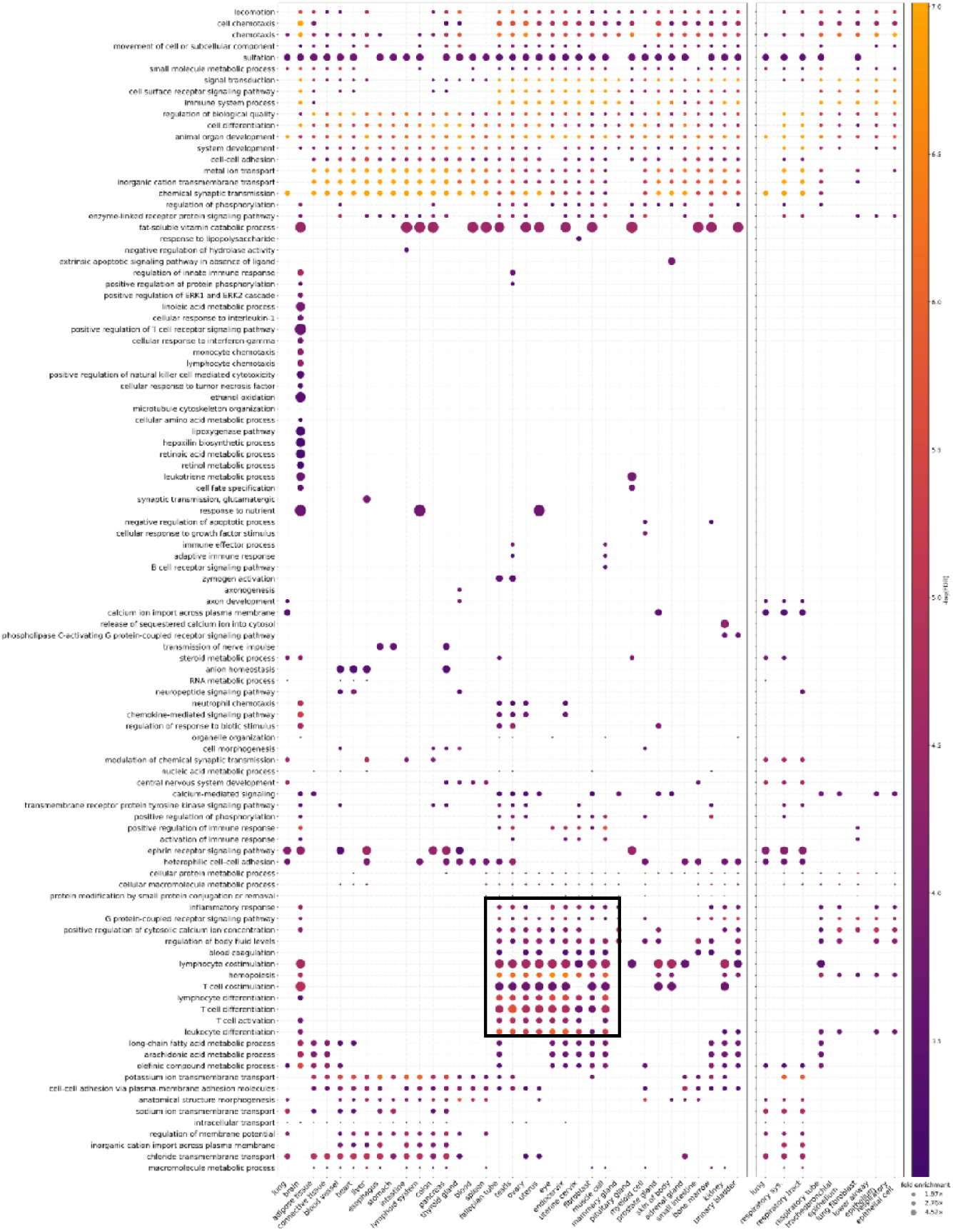
Mahi models context-specific gene perturbations. Enriched biological processes for all displayed tissues. Lung and related cell-types and tissues have similar enrichment signals compared to other contexts. Reproductive tissues are also enriched for a similar set of distinctive pathways, including immune response and CFTR-dependent epithelial hydration.

